# Square beams for optimal tiling in TEM

**DOI:** 10.1101/2023.07.29.551095

**Authors:** Eugene YD Chua, Lambertus M Alink, Mykhailo Kopylov, Jake Johnston, Fabian Eisenstein, Alex de Marco

**Affiliations:** Simons Electron Microscopy Center, New York Structural Biology Center, New York, NY 10027; Department of Physiology and Cellular Biophysics, Columbia University, New York, NY, USA; Graduate School of Medicine, University of Tokyo, Tokyo, Japan

**Keywords:** TEM, cryo-EM, square beam, TEM tiling

## Abstract

Imaging large fields of view at a high magnification requires tiling. Transmission electron microscopes typically have round beam profiles; therefore, tiling across a large area is either imperfect or results in uneven exposures, a problem on dose-sensitive samples. Here, we introduce a square electron beam that can be easily retrofitted in existing microscopes and demonstrate its application, showing it can tile nearly perfectly and deliver cryo-EM imaging with a resolution comparable to conventional setups.

## Main text

In transmission electron microscopy (TEM) of dose-sensitive specimens such as vitrified biological material, pre-exposure of areas to the beam attenuates the attainable resolution (Baker & Rubinstein, 2010). High-resolution cryo-TEM typically comes at the cost of a reduced field of view; therefore, a balance between the pixel size and the sample imaged within its biological context is required. Since the illumination profile (which we call here “beam” or “beam profile” for simplicity) of TEMs is round, tiling across a large field of view encounters the circle packing problem, wherein circles cannot be perfectly tiled. Even with an ideal modern imaging setup with a fringe-free imaging (FFI) (Konings et al., 2019; Weis & Hagen, 2020) and a square sensor, the sensor will only capture ∼69% of the area illuminated by the tightest possible round beam (Figure 1 and Supplementary Figure S1). The electron beam will damage the remaining illuminated but unimaged site, which will no longer contain high-resolution information when next imaged. This is a well-known limitation in montage tomography of vitrified specimens, and while data collection schemes that account for overlapping exposures exist (Peck et al., 2022; Yang et al., 2022), the illumination across multiple exposed areas remains non-uniform.

**Figure 1.**
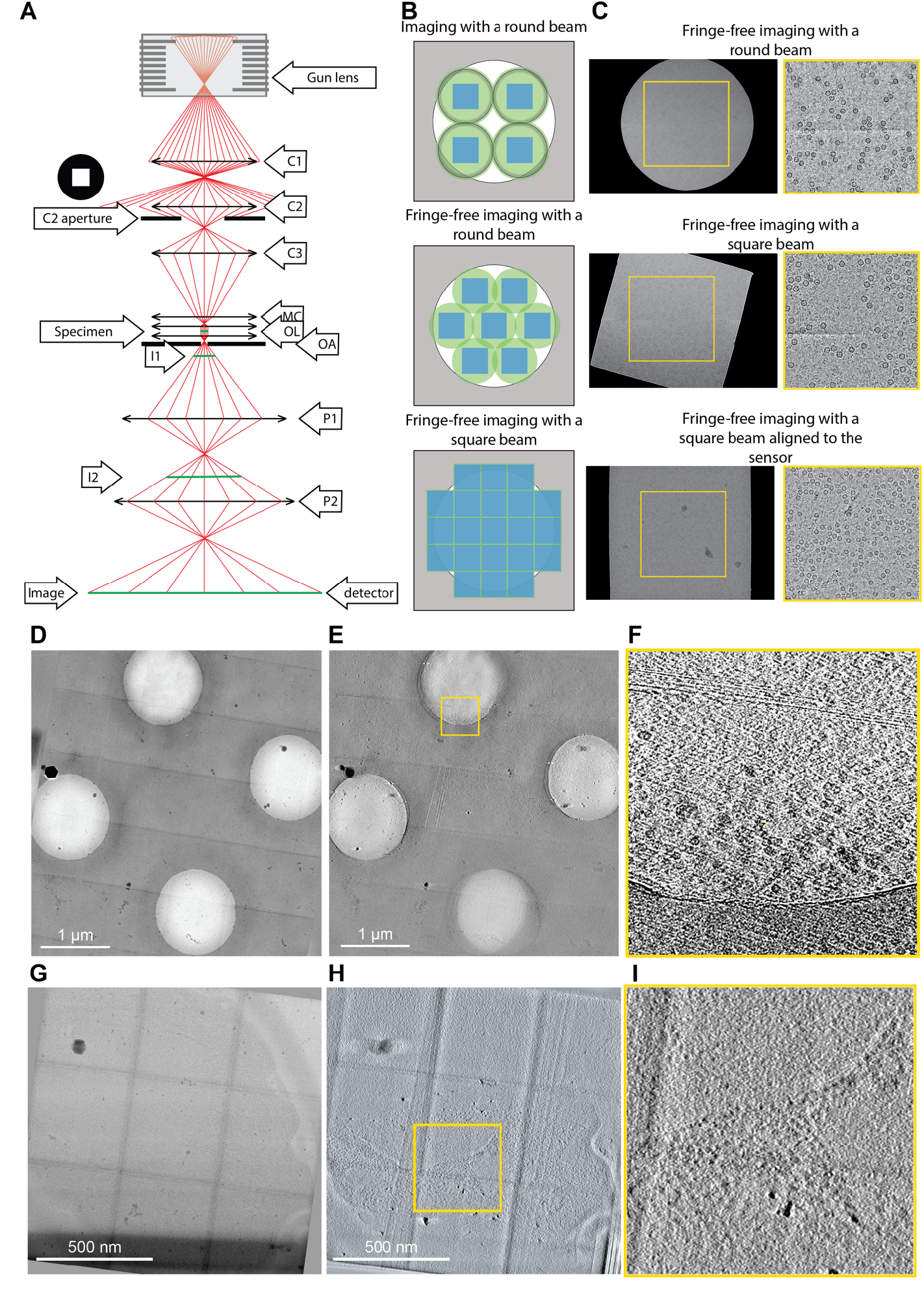
(A) Ray diagram of electrons passing through the column of a TEM. The square aperture is placed at the “C2 aperture” position. C1/2 - condenser lens 1/2, OL - objective lenses, OA - objective aperture, MC - minicondenser, I1/2 - intermediate image plane 1/2, P1/2 - projection lens 1/2. (B) Examples of imaging a specimen (white circle) with different TEM beam setups. The areas on the specimen illuminated by the electron beam are shown in green circles (round beam) or green squares (square beam); the illuminated areas are top image, black rings) in non-fringe-free TEM setups, the beam must be spread out so the fringes do not fall on the sensor. (C) Example micrographs from an FFI-enabled TEM, acquired with a round beam (top), square beam (middle), and a square beam aligned to the sensor by adjusting the projection lens (bottom). (D) Example of PACE-tomo tiled imaging with the square beam. We collected a 5x5 butt-joint tile set on holey carbon grids with apoferritin. (E) The resulting tomogram reconstructed from the data shown in D. (F) High magnification crop of the joint between two tiles from panel E. In the upper section, imperfections in the stitching are visible, as no alignment or interpolation was performed when stitching. However, the apoferritin particles are clearly visible. (G) PACE-tomo tiled imaging with the square beam on yeast lamellae. Here, we collected a 3x3 butt-joint tile set. (H) The resulting tomogram reconstructed from the data shown in G. (I) High magnification crop of a region of the tomogram.

One solution to the problem of imperfect tiling with round beams is to use a square electron beam. Modern cryo-TEMs use Mueller-type sources where the emitter’s shape defines the electron beam shape, typically resulting in a circle. For TEM imaging in a 3-condenser system, the beam current (spot size) is selected by the C1 and C2 lenses, and the source beam width (beam convergence) is determined by changing the strength of the C2 and C3 lenses. The aperture between the C2 and C3 lenses becomes the beam-shaping aperture. When the electron beam cross-over above the C2 aperture is moved (by changing C1/C2 lenses), the beam current changes, whereas when the cross-over below the C2 is forced (by changing the C2/C3 lenses), the size of the beam changes. The post-C2-aperture beam takes on the shape of the aperture’s hole when the beam is spread more comprehensively than the aperture. For practical reasons linked to manufacturing and isotropic optical propagation, all cracks have round holes, creating round beams. In this work, we use a C2 aperture with a square hole to create a square electron beam profile. We demonstrate its utility on an FFI-capable TEM for near-perfect tiling in montage tomography and increased efficiency in data collection for single particle analysis with minimal loss of resolution.

Using a square C2 aperture, we successfully created a square beam (Figure 1C). First, we adjusted the beam width using the microscope intensity control such that the beam had the same size as the shortest dimension of the sensor. Then, the (post-objective) projection P2 lens was adjusted to rotate the beam square onto the sensor (Figure 1C). Aligning the beam with the sensor ensures that the sensor images the entire sample area exposed to the beam. Since changes in the P2 lens strength changed the image’s rotation, magnification, and defocus, calibrations for pixel size, image shift, and eucentric focus had to be redone. The flux on the sensor can be adjusted with spot size, and the beam intensity distribution across the illuminated area can be measured to ensure uniform exposures (**Error! Reference source not found**.). The unique P2 lens state can be stored as a unique magnification entry and added as a separate registry key.

With a square beam, it became possible to tile with minimal overlap to exhaustively image a large field of view. This is especially important for *in situ* tomography, where it is often helpful to image large contiguous areas of a specimen, such as a lamella, at high resolution. Acquisition targets can be set along the tilt axis to overlap minimally and, therefore, reduce any areas on the sample that are exposed to the electron beam in more than one acquisition target. To maximize the acquisition area while avoiding the sample overexposure in the direction perpendicular to the tilt axis, a new data acquisition scheme was developed, where the beam-image shift is independent of the sample tilt. This scheme was implemented in PACE-tomo (Eisenstein et al., 2023), and the beam shifts were set to be equivalent to one image Y-axis in nm (Figure 1D-I and Supplementary Videos). The beam overlap was maintained identically throughout the tilt series (shows the difference between conventional tiled at 0 deg versus camera-based offset). After data acquisition, a montage for each stage tilt can be stitched to produce a sizeable field-of-view image, which is then aligned across the tilt series and reconstructed.

In single particle data acquisition, a square beam significantly increases throughput. Using common supports such as UltrAuFoil R1.2/1.3 grids, a square beam could image up to 5 targets per hole (85 targets per stage movement) versus 2 targets per hole for a round beam (Supplementary Figure S4). This increased the data collection rate by nearly 2-fold, from ∼158 exposures per hour with the round beam to ∼291 exposures per hour with the square beam.

While we observed normal behavior during microscope alignment and coma correction (Supplementary Figure S5), we consistently obtained a slightly worse reconstruction B-factor for the square beam. However, with enough particles, the reconstruction still went to Nyquist (Figure 2). With smaller particle sets, we consistently observed a slight loss in reconstruction resolution (∼0.1 Å) with the square beam compared to the round beam (Supplementary Figure S8). This is most likely linked to the lack of circular symmetry in the phase profile of the beam diffraction from the square aperture. A potential solution is to use larger apertures to lower the diffraction angle and maintain a more uniform phase profile at the sample plane.

**Figure 2.**
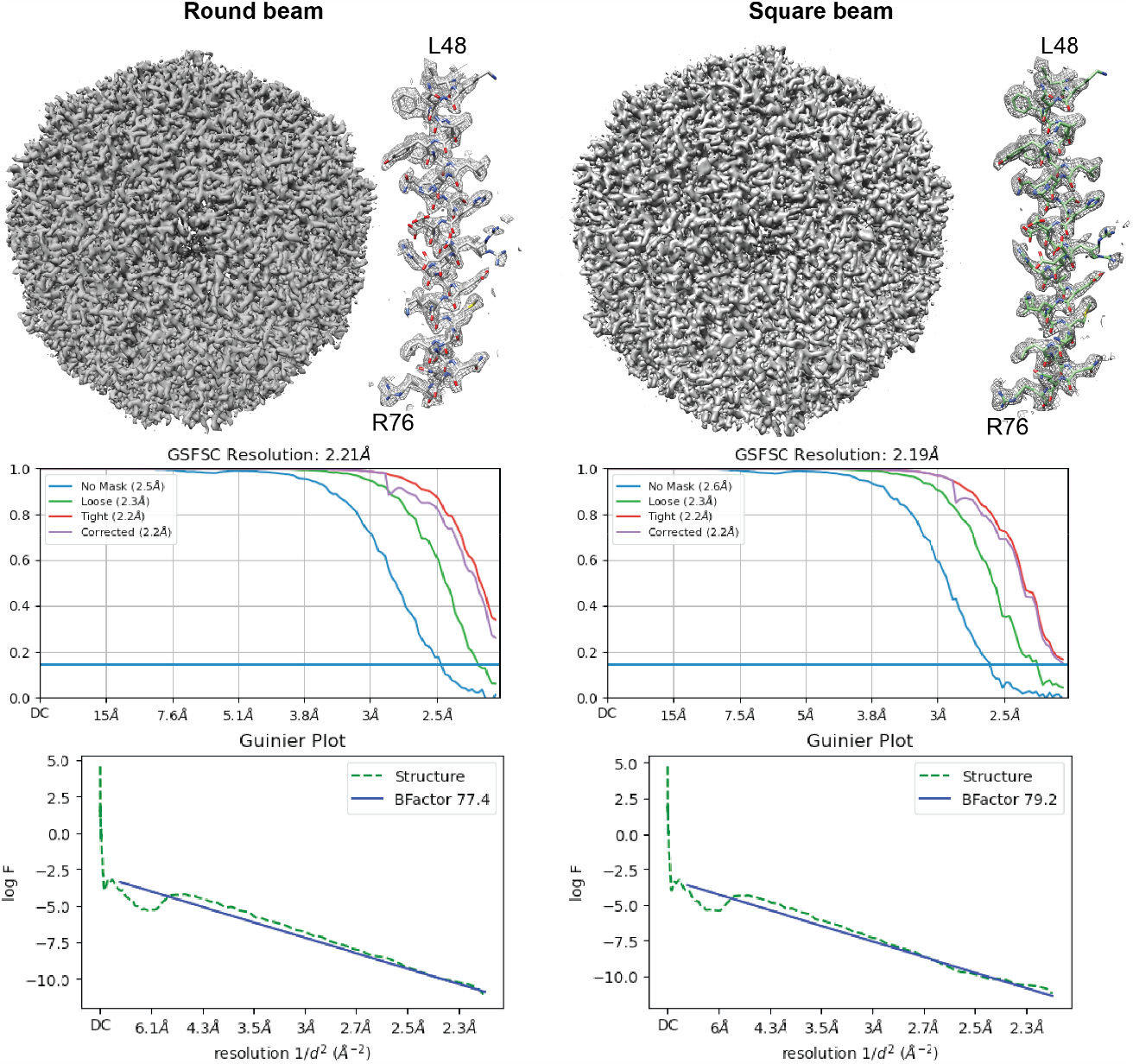
Single particle reconstructions of apoferritin with data collected with a round (left) or a square (right) beam. In both cases, with 120,000 particles, the reconstructions can achieve Nyquist resolution.

The most notable change from the round to the square beam is that the spherical aberration fit increases from 2.9 to 3.1 mm during homogeneous refinement. The increased spherical aberration happens with the detuning of the P2 lens – the spherical aberration did not change with square aperture data collected without P2 lens detuning.

Considering the change in the tuning of the P2 lens, we tested the potential changes in microscope performance. The magnification was not subject to any measurable distortion (Supplementary Table S1) (Grant & Grigorieff, 2015). Further, Young’s Fringes experiment showed that the microscope performance remained within the manufacturer’s specification as the information transmittance reached 0.14 nm on gold cross-grating (Supplementary Figure S9).

Non-circular and square beams have been developed for beam-shaping in electron beam lithography, laser micromachining, and medical laser applications. We use a square beam for nearly perfect cryo-EM and cryo-ET montage tiling. Other non-round beams, such as rectangles and hexagons, which are also optimal profiles for tiling, can be used (Brown et al., 2023). We note that it is possible to create a rectangular beam by stigmating a square beam; however, this introduces unwanted aberrations into the beam. Furthermore, matching the beam’s and detector’s shapes prevents minor data processing complications associated with having unilluminated sensor areas.

Among optimizations, the alignment of the aperture orientation with respect to the sensor is critical to ensure optimal overlap between the sensor and the illuminated area. As an alternative to rotating the beam with the P2 lens, we envision that the aperture can be mechanically rotated through a redesign of the aperture strip: a worm wheel gear can be installed to physically rotate the square aperture while it is in the liner tube under vacuum during illumination. For this work, we propose a no-cost solution that involves adjusting the P2 lens’ current to induce a rotation of the projected image plane. This introduces changes in the optical system, affecting the image’s magnification, defocus, and rotation, requiring recalibration of the pixel size, eucentric focus, and image-shift matrices.

## Acknowledgments

We thank Dr. Masahide Kikkawa (University of Tokyo) for the apoferritin plasmid and Dr. Brian Kloss (NYSBC) for expressing and purifying the protein. We thank Anchi Cheng, Bridget Carragher, and Clint Potter for the early discussions and support.

This work was supported by the Simons Electron Microscopy Center and National Resource for Automated Molecular Microscopy located at the New York Structural Biology Center, supported by grants from the Simons Foundation (SF349247) and the NIH National Institute of General Medical Sciences (GM103310).

## Data availability

Data will be made freely available upon request.

## Supplementary material

### Methods

Square apertures were purchased from Agar Scientific (product numbers AGAS3005P and AGAS3005P). The apertures are made of platinum, with a diameter of 3.04 mm, a thickness of 0.25 mm, and a square hole of 50 or 100 µm. A 50-micron aperture has been installed into the C2 aperture holder of the microscope. Before its installation into the microscope, the square aperture should be plasma cleaned to remove any impurities, then maintained in a sealed container for several days to allow the charge to dissipate, allowing an easier insertion into the aperture strip.

The optics of the electron microscope consists roughly of three sections: the beam forming condenser section where the square aperture is installed, the objective section with the specimen, and the magnifying projection lens section. See also Figure 1A. The square aperture is located at the condenser section, defining the square beam. To align the square beam towards the camera, we tuned the P2 projection lens. The condenser and objective lens settings were not touched to keep the parallelism of the beam intact going through the objective system. With the electron microscope set to diffraction mode, on de-tuning of the condenser lens, the diffraction rings showed blurring, indicating loss of parallelism of the beam in the objective lens. Detuning of the projection system left the parallelism intact. This was expected since the projection system is below the objective lens. Note that tuning the projection lens will affect the image’s magnification, rotation, and defocus.

The strength of the P2 lens can be checked in the user interface system status overview (Supplementary Figure S6). The square aperture can be inserted using the standard OCX in the user interface. The square C2 aperture has the same form factor as the ‘standard’ round C2 aperture. Replacement of the C2 aperture, aligning the aperture laterally, and tuning the P2 lens can be done by the equipment supplier service engineer using the standard software and wizards. A screenshot of the aperture wizard is given in Supplementary Figure S7. After the adjustments, eucentric focus calibration, pixel size, and image shift calibrations must be performed in the SerialEM (Mastronarde, 2005).

The same protein sample was used for apoferritin tomography and single particle analysis prepared as described below, with the only difference found in the support (carbon vs gold). Data were acquired using PACE-tomo scripts (Eisenstein et al., 2023) incorporated into SerialEM 4.1 beta 13 (Mastronarde, 2005) with a pixel size of 2.12 Å/px, exposure dose of 3.4 e^-^/Å^2^ per tilt, and -45° to 45° tilt range for a total of 31 tilts and total dose of 105 e^-^/Å^2^ per tilt series. Imaging was done as 5x5 patches. Saccharomyces cerevisiae standard yeast test sample for lamella tomography was prepared following the Waffle method protocol (Kelley et al., 2022; Klykov et al., 2022). Data were acquired using PACE-tomo scripts incorporated into SerialEM 4.1 beta 13 with a pixel size of 2.12 Å/px, exposure dose of 2.55 e^-^/ Å^2,^ and a total dose of 76.5 e^-^/ Å^2^. Imaging was done as 3x3 patches to encompass the whole yeast cell in the field of view.

For the apoferritin dataset, the acquired tilt series were first motion corrected with Warp (Tegunov & Cramer, 2019), and tilt series were aligned and reconstructed with AreTomo (Zheng et al., 2022) at bin6 and used without additional processing for further analysis. For the yeast lamella dataset, the tilt series were motion-corrected with Warp (Tegunov & Cramer, 2019) and aligned using AreTomo (Zheng et al., 2022). Tomogram reconstruction was performed using Tomo3D (Agulleiro & Fernandez, 2015), and the tomogram was deconvolved IsoNet (Liu et al., 2022) to enhance contrast. Stitching for both the apoferritin and yeast lamella datasets was done automatically with custom Python scripts, where the tiles were stitched together using the image shift locations obtained from the metadata.

For single particle analysis, UltrAuFoil R1.2/1.3 300 mesh Au grids were hydrophilized with a mixture of Ar and O_2_ gas (26.3:8.7 ratio) at 15 W for 7 seconds in a Solarus Model 950 Advanced Plasma System (Gatan). 3 µl of 8 mg/ml mouse apoferritin was pipetted onto each grid, blotted for 3-5 seconds in a Vitrobot at 20°C and 100% relative humidity, then vitrified in liquid ethane. The P2 projection lens was detuned to rotate the square beam square onto the sensor, resulting in changes in the image’s magnification, rotation, and defocus. Eucentric focus needed to be reset on the microscope by adjusting the objective lens, then beam and image shift and scale rotation calibrations needed to be redone in Leginon (Cheng et al., 2021; Suloway et al., 2005) prior to data collection. Pixel size calibration was done in SerialEM on a standard cross-grating replica grid. Energy filter alignments were done per standard protocol, with the entire sensor illuminated. Objective lens astigmatism and coma correction were performed using Sherpa, with the full sensor illuminated. Single particle data were collected using Leginon with either a 100 µm round C2 aperture or a 50 µm square C2 aperture. Data were collected at a pixel size of ∼1.08 Å/pixel, a flux of ∼30 e/px/s for 2 seconds, equaling a total dose of ∼51 e/Å^2^, with a nominal defocus range of -0.5 µm to -2.0 µm. The square beam was condensed to match the size of the sensor to maximize the data acquisition area, and in the control experiment, a round beam was used with its intensity set to match the flux of the square beam on the sensor. Data were collected using a beam-image shift (Cheng et al., 2018).

For single particle data processing, square and round beam data was first motion corrected with patches in cryoSPARC v4.2.1 (Punjani et al., 2017). Full micrographs were then patch CTF estimated, and particles were picked using apoferritin templates. Particles were extracted with a box size of 280 pixels, 2D classified, and 120,000 particles randomly selected for homogeneous refinement. To exclude the unilluminated areas of the sensor because of the condensed square beam, a central square region of the motion-corrected micrographs was cropped out with the IMOD (Mastronarde & Held, 2017) command “trimvol”. As a control, the same central square region was cropped from both square and round aperture data. Cropped micrographs were then re-imported into cryoSPARC, and the same processing workflow continued as above. Reconstructions from full and cropped micrographs from square and round aperture data were compared.

**Supplementary Figure S1.**
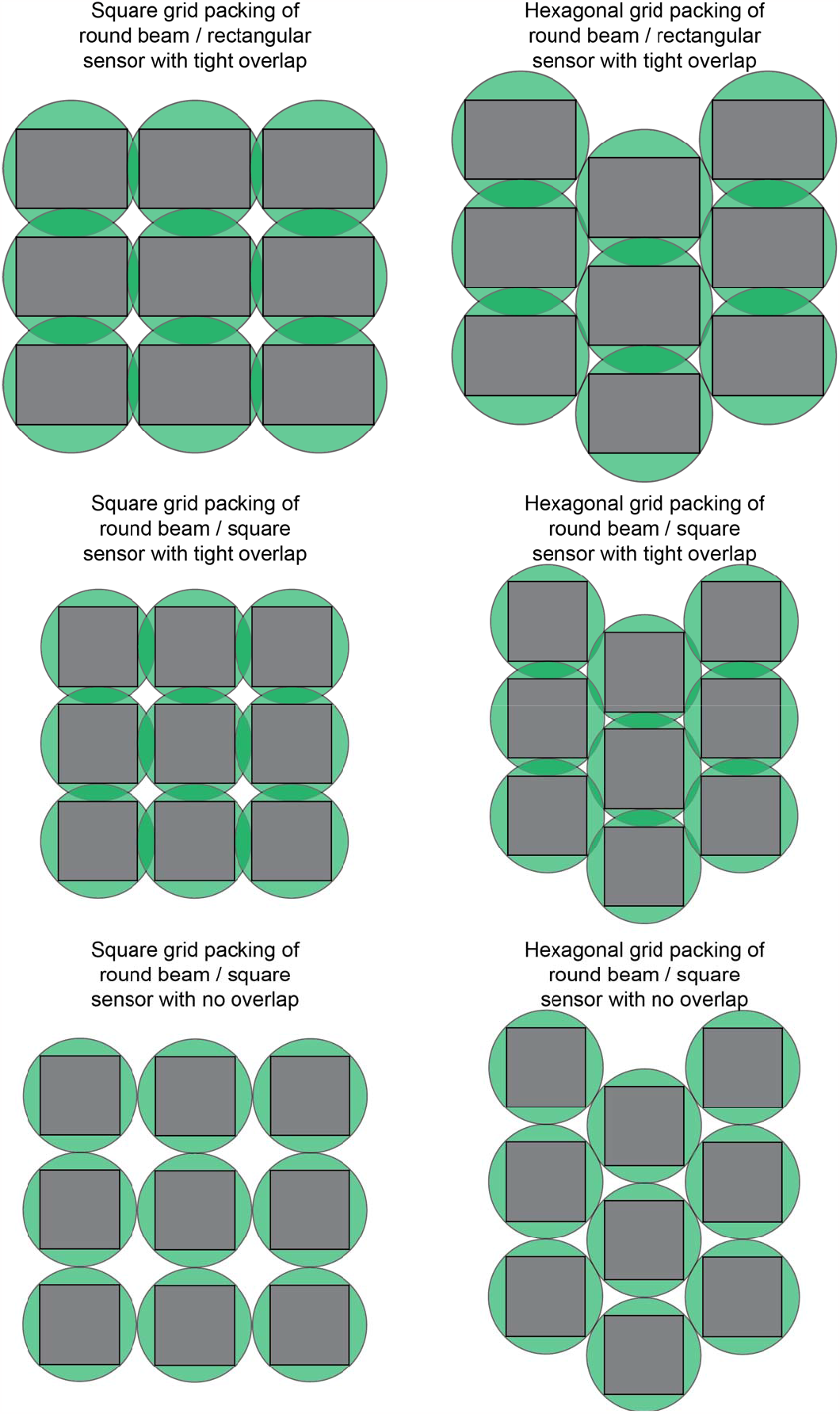
Examples of packing circular beams (green) and square or rectangular sensors (grey) highlighting the gap between the exposed and imaged areas.

**Supplementary Figure S2.**
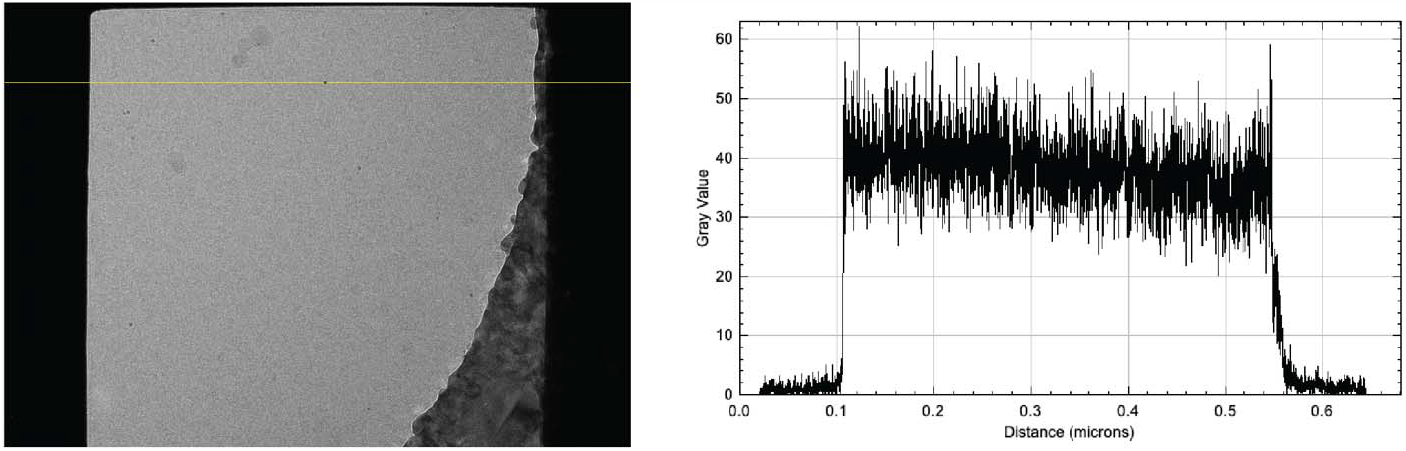
Example micrograph taken with a square beam (left), showing a pixel intensity profile (right) for the pixels along the yellow line. Pixel intensity profile was obtained with ImageJ (Schneider et al., 2012).

**Supplementary Figure S3.**
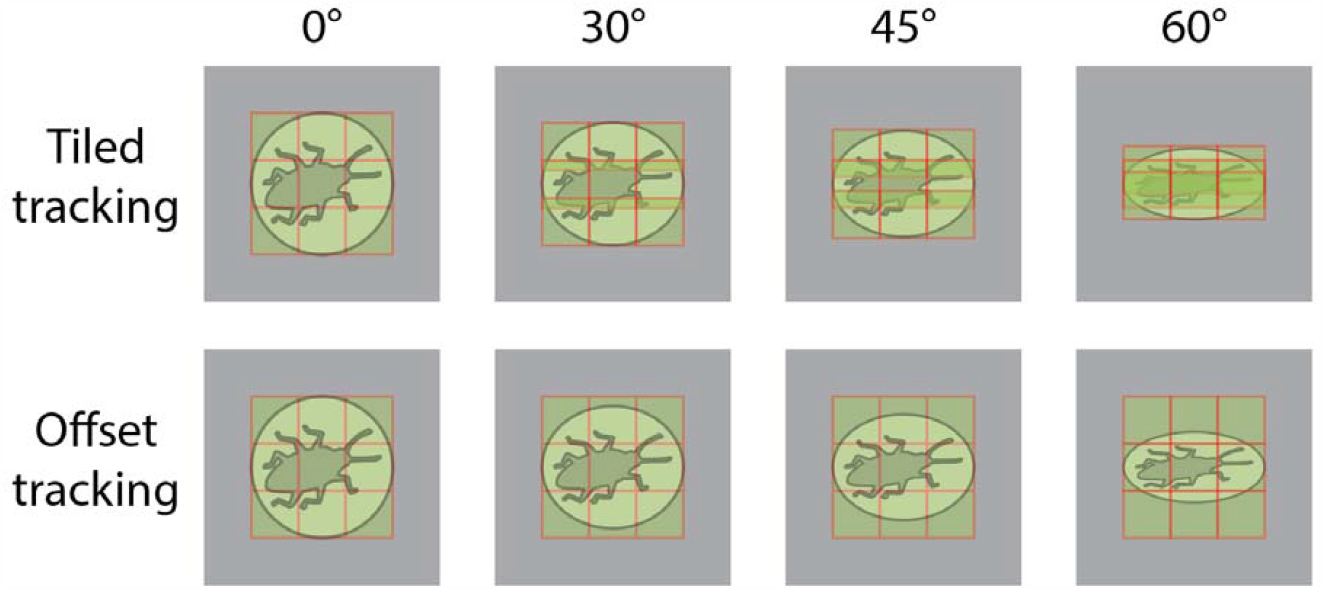
Tomography data acquisition schemes with the square beam and the effect of sample tilt on beam overlap when using a fixed beam offset (tiled tracking). We implemented an acquisition scheme where the position of the tile is determined by the Y-axis length of the image in nm (offset tracking).

**Supplementary Figure S4.**
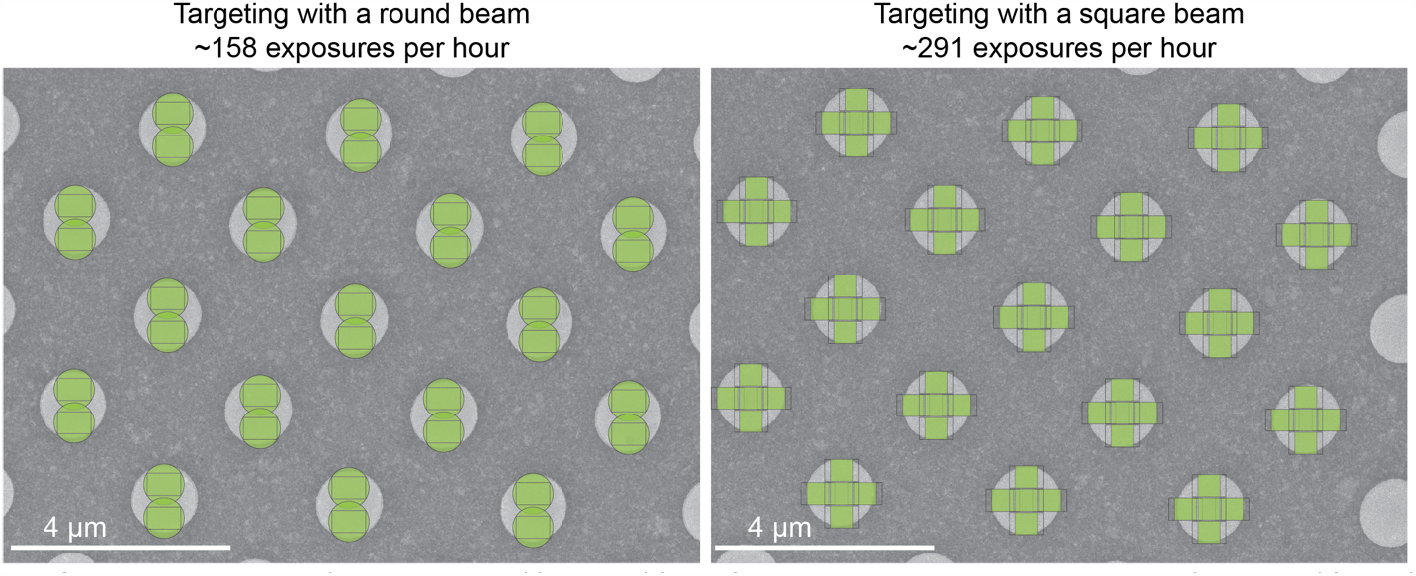
When using a round beam and fringe-free imaging, two acquisition targets can be acquired for each 1.2 µm hole (left). In this field of view, up to 34 acquisition images can be taken per stage movement. When using a square beam with perfect tiling, five acquisition targets can be acquired for each 1.2 µm hole (right), increasing the number of images to 85 per stage movement.

**Supplementary Figure S5.**
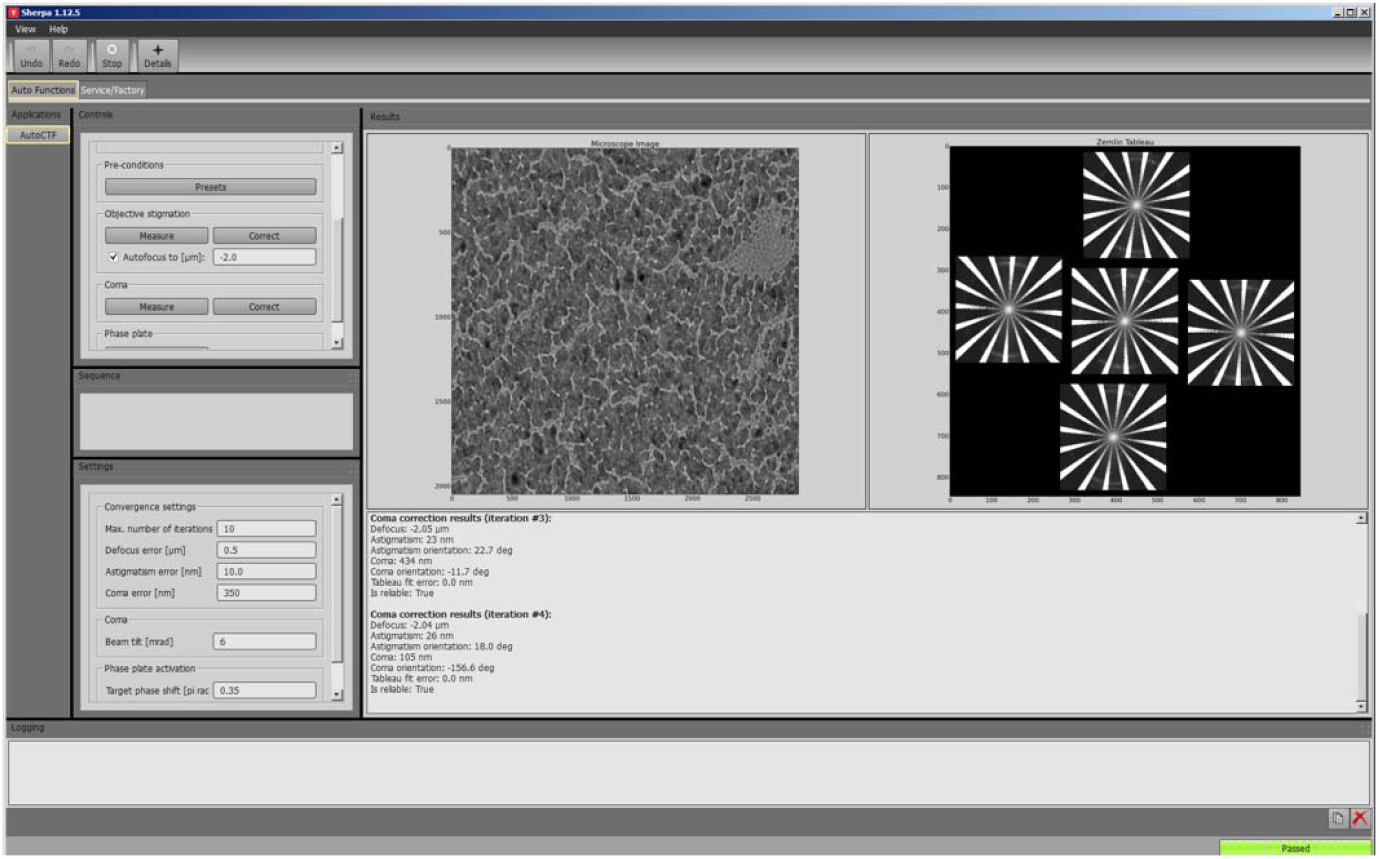
Automated coma correction while using a square beam shows it is possible to achieve acceptable levels of coma, directly comparable to using round illumination.

**Supplementary Figure S6.**
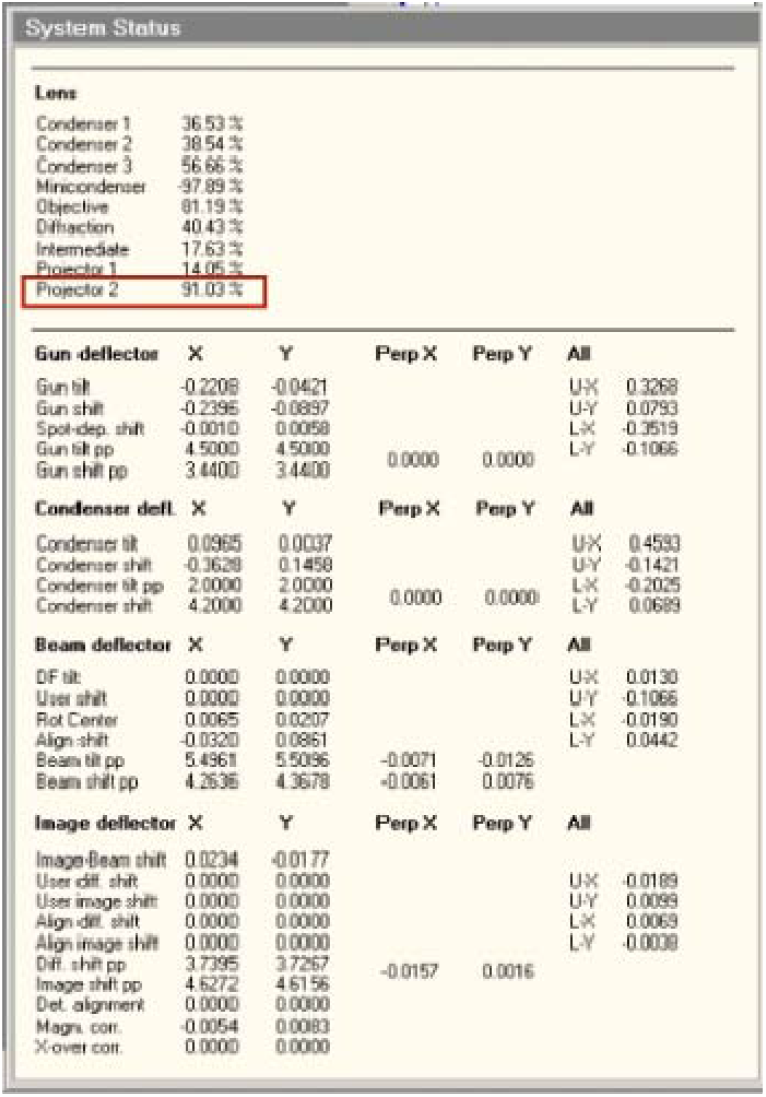
System status overview in the user interface showing the strength of the P2 lens.

**Supplementary Figure S7.**
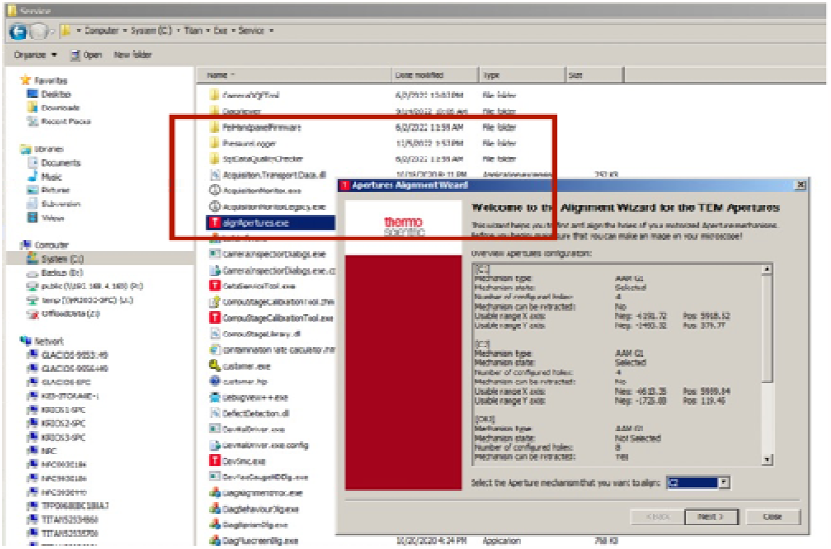
Aperture alignment wizard that allows adjusting the C2 aperture.

**Supplementary Figure S8.**
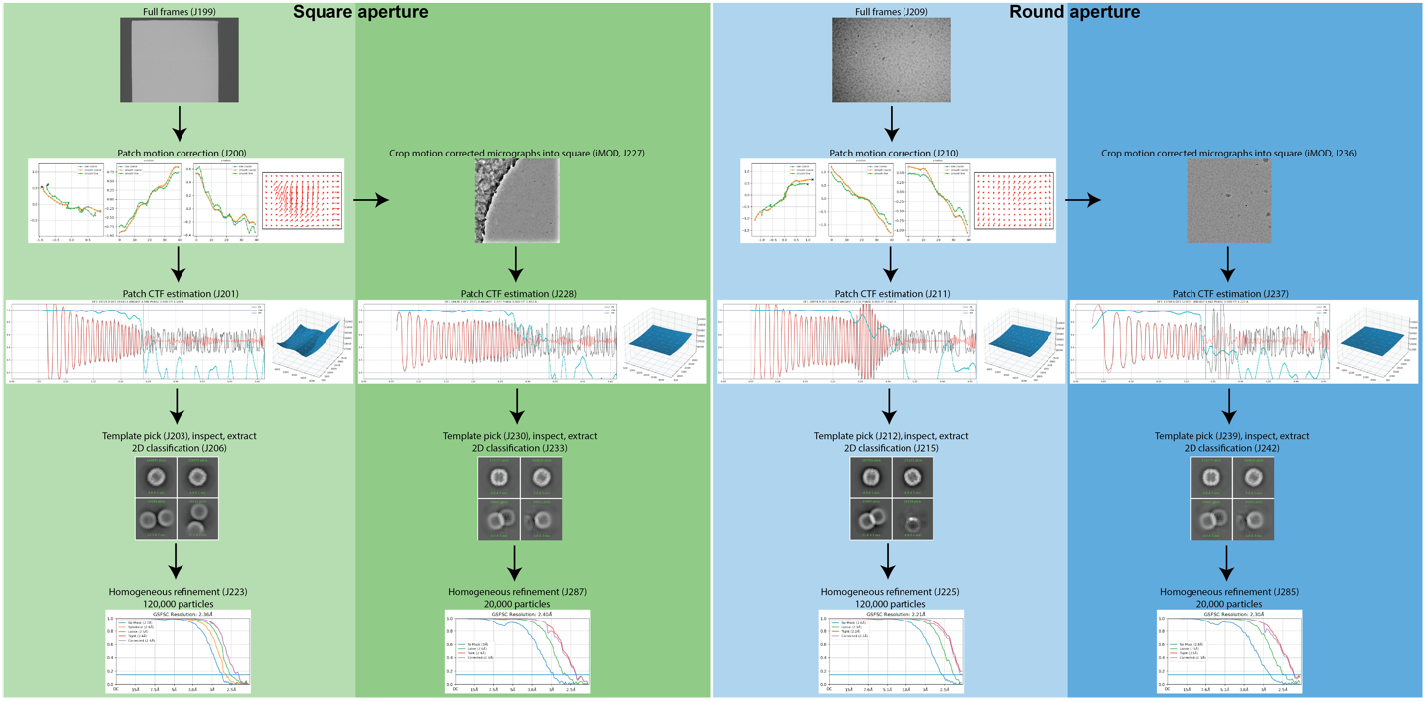
Single particle data processing workflow for the square (green) and round (blue) apertures. For each aperture type, the processing workflow uses the full frames on the left, and the processing workflow with cropped micrographs (to exclude the unilluminated areas of the sensor) on the right.

**Supplementary Table S1.** Anisotropic magnification distortion. We performed a common measurement of the magnification along all axes using the program described by Grant and Grigorieff (Grant & Grigorieff, 2015). We tested two different apertures and imaged with the conventional (P2 lens rotated? = No) and new P2 lens tuning (P2 lens rotated? = Yes). All the conditions show the same distortion.

| C2 aperture | P2 lens rotated? | Magnification | Distortion angle (degrees) | Major axis scale factor | Minor axis scale factor | Distortion (%) |
| --- | --- | --- | --- | --- | --- | --- |
| Round, 100 $\mu\text{m}$ | No | 81,000 | 8.6 | 1.002 | 0.998 | 0.45 |
| Square, 50 $\mu\text{m}$ | No | 81,000 | 10.5 | 1.002 | 0.998 | 0.45 |
| Round, 100 $\mu\text{m}$ | Yes | ~81,000 | 27.6 | 1.002 | 0.998 | 0.35 |

**Supplementary Figure S9.**
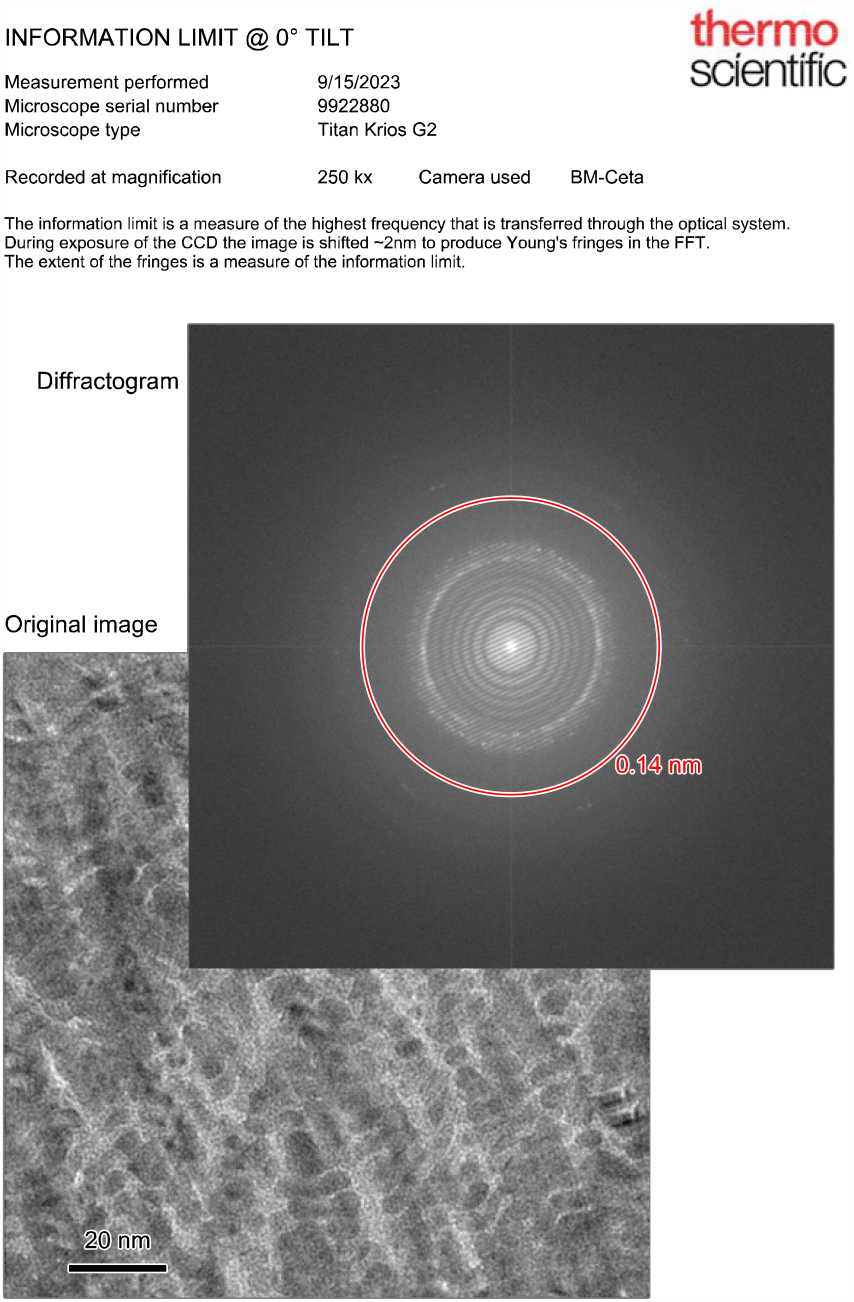
Screenshot of the standard output from the Young’s fringes test performed by Thermo Fisher service. This test shows the transmittance limit of the microscope, and in this case, it demonstrates that the use of a square aperture and a differently tuned projection system does not impact on microscope performance.

## References

Agulleiro, J.-I., & Fernandez, J.-J. (2015). Tomo3D 2.0 – Exploitation of Advanced Vector eXtensions (AVX) for 3D reconstruction. Journal of Structural Biology, 189(2), 147–152. 10.1016/j.jsb.2014.11.009

Baker, L. A., & Rubinstein, J. L. (2010). Radiation Damage in Electron Cryomicroscopy. In Methods in Enzymology(Vol. 481, pp. 371–388). Elsevier. 10.1016/S0076-6879(10)81015-8

Brown, H. G., Smith, D., Wardle, B. C., & Hanssen, E. (2023). Fitting a square beam in a square camera: Novel condenser apertures for low-dose transmission electron microscopy [Preprint]. Biophysics. 10.1101/2023.08.13.553155

Cheng, A., Eng, E. T., Alink, L., Rice, W. J., Jordan, K. D., Kim, L. Y., Potter, C. S., & Carragher, B. (2018). High resolution single particle cryo-electron microscopy using beam-image shift. Journal of Structural Biology, 204(2), 270–275. 10.1016/j.jsb.2018.07.015

Cheng, A., Negro, C., Bruhn, J. F., Rice, W. J., Dallakyan, S., Eng, E. T., Waterman, D. G., Potter, C. S., & Carragher, B. (2021). Leginon: New features and applications. Protein Science, 30(1), 136–150. 10.1002/pro.3967

Eisenstein, F., Yanagisawa, H., Kashihara, H., Kikkawa, M., Tsukita, S., & Danev, R. (2023). Parallel cryo electron tomography on in situ lamellae. Nature Methods, 20(1), 131–138. 10.1038/s41592-022-01690-1

Grant, T., & Grigorieff, N. (2015). Automatic estimation and correction of anisotropic magnification distortion in electron microscopes. Journal of Structural Biology, 192(2), 204–208. 10.1016/j.jsb.2015.08.006

Kelley, K., Raczkowski, A. M., Klykov, O., Jaroenlak, P., Bobe, D., Kopylov, M., Eng, E. T., Bhabha, G., Potter, C. S., Carragher, B., & Noble, A. J. (2022). Waffle Method: A general and flexible approach for improving throughput in FIB-milling. Nature Communications, 13(1), 1857. 10.1038/s41467-022-29501-3

Klykov, O., Bobe, D., Paraan, M., Johnston, J., Potter, C., Carragher, B., Kopylov, M., & Noble, A. (2022). In situ cryo-FIB/SEM Specimen Preparation Using the Waffle Method. BIO-PROTOCOL, 12(21). 10.21769/BioProtoc.4544

Konings, S., Kuijper, M., Keizer, J., Grollios, F., Spanjer, T., & Tiemeijer, P. (2019). Advances in Single Particle Analysis Data Acquisition. Microscopy and Microanalysis, 25(S2), 1012–1013. 10.1017/S1431927619005798

Liu, Y.-T., Zhang, H., Wang, H., Tao, C.-L., Bi, G.-Q., & Zhou, Z. H. (2022). Isotropic reconstruction for electron tomography with deep learning. Nature Communications, 13(1), 6482. 10.1038/s41467-022-33957-8

Mastronarde, D. N. (2005). Automated electron microscope tomography using robust prediction of specimen movements. Journal of Structural Biology, 152(1), 36–51. 10.1016/j.jsb.2005.07.007

Mastronarde, D. N., & Held, S. R. (2017). Automated tilt series alignment and tomographic reconstruction in IMOD. Journal of Structural Biology, 197(2), 102–113. 10.1016/j.jsb.2016.07.011

Peck, A., Carter, S. D., Mai, H., Chen, S., Burt, A., & Jensen, G. J. (2022). Montage electron tomography of vitrified specimens. Journal of Structural Biology, 214(2), 107860. 10.1016/j.jsb.2022.107860

Punjani, A., Rubinstein, J. L., Fleet, D. J., & Brubaker, M. A. (2017). cryoSPARC: algorithms for rapid unsupervised cryo-EM structure determination. Nature Methods, 14(3), 290–297. 10.1038/nmeth.4169

Schneider, C. A., Rasband, W. S., & Eliceiri, K. W. (2012). NIH Image to ImageJ: 25 years of image analysis. Nature Methods, 9(7), 671–675. 10.1038/nmeth.2089

Suloway, C., Pulokas, J., Fellmann, D., Cheng, A., Guerra, F., Quispe, J., Stagg, S., Potter, C. S., & Carragher, B. (2005). Automated molecular microscopy: The new Leginon system. Journal of Structural Biology, 151(1), 41–60. 10.1016/j.jsb.2005.03.010

Tegunov, D., & Cramer, P. (2019). Real-time cryo-electron microscopy data preprocessing with Warp. Nature Methods, 16(11), 1146–1152. 10.1038/s41592-019-0580-y

Weis, F., & Hagen, W. J. H. (2020). Combining high throughput and high quality for cryo-electron microscopy data collection. Acta Crystallographica Section D Structural Biology, 76(8), 724–728. 10.1107/S2059798320008347

Yang, J. E., Larson, M. R., Sibert, B. S., Kim, J. Y., Parrell, D., Sanchez, J. C., Pappas, V., Kumar, A., Cai, K., Thompson, K., & Wright, E. R. (2022). Correlative cryogenic montage electron tomography for comprehensive in-situ whole-cell structural studies. bioRxiv.10.1101/2021.12.31.474669

Zheng, S., Wolff, G., Greenan, G., Chen, Z., Faas, F. G. A., Bárcena, M., Koster, A. J., Cheng, Y., & Agard, D. A. (2022). AreTomo: An integrated software package for automated marker-free, motion-corrected cryo-electron tomographic alignment and reconstruction. Journal of Structural Biology: X, 6,100068. 10.1016/j.yjsbx.2022.100068

